# First confirmed record in Melilla of *Gerbillus campestris* Loche, 1867 (Muridae: Gerbillinae) for Spain

**DOI:** 10.64898/2026.09.30.754369

**Authors:** Trino Ferrández Verdú, Francisco Pérez Ruiz, Antonio Félix Carrillo López, Manuel Tapia Claro, Alicia Castillo González, Rubén Vives López, Claudio Amorós Box, Mario Marcos Pérez, Zbyszek Boratyński, Barbora Pavlíčková

## Abstract

The fieldwork was carried out between December 2023 and June 2025, with the aim, among others, of confirming the possible presence of the North African Gerbil (*Gerbillus campestris*) in Melilla. To this end, 56 camera trap stations were installed, each equipped with a camera trap. Of the total number of camera trap stations installed, the presence of *Gerbillus campestris* was detected in six of them, representing a capture success rate of 10.71%. Following the detection of the species, a specimen was captured using cage traps. For identification purposes, samples of epithelial tissue from the ear and faeces were collected. These samples were sent to the Institute of Vertebrate Biology of the Czech Academy of Sciences, Brno, Czech Republic, for analysis. There, DNA was extracted from the samples using the Innu PREP DNA Mini Kit 2.0 (IST Innuscreen GmbH, Germany) according to the manufacturer’s protocol. The results obtained confirm, for the first time through records of the species, the presence of Gerbillus campestris in Melilla and, therefore, in Spain and Europe.

## Introduction

*Gerbillus campestris* Loche, 1867, the North African Gerbil, belongs to the subgenus *Dipodillus* from the genus *Gerbillus*. It occurs in North Africa from Morocco to Egypt, with a disjunct distribution in the Sahara and in sub-Saharan Africa, including Mali, Chad, Niger, and Sudan (Musser and Carleton, 2005). This nocturnal gerbil is a small yellow to brown gerbil with a long tail (130-140 % of head-body length). It has bare soles on its hindfeet. Its ventral pelage is white with strict delineations from flanks, and the tail is bicolored with a pencil on the terminal end. This gerbil inhabits areas near agricultural lands, such as cultivated fields, as well as rocky desert environments, including inland cliffs and mountain peaks. It is considered an agricultural pest in North Africa. The Red List of IUCN classifies it as Least Concern (Granjon, 2016; Wilson et al. 2016).

## Materials and Methods

### Study Area

Melilla, the city where the fieldwork was carried out, is a heavily human-modified territory with a surface area of 12.3 km^2^, yet it is situated within a unique biogeographical context. The city is located on the eastern coast of the Guelaya Peninsula, which extends into the Alboran Sea, its tip being Cape Tres Forcas. This peninsula has a complex topography and high ecological value. The city is also very close to the Mar Chica lagoon (to the southeast) and Mount Gurugú (to the southwest), other unique areas of environmental value, already within Moroccan territory.

Approximately 55% of Melilla’s territory is occupied by significantly transformed areas: residential, infrastructural, industrial, military, etc. The remaining 45%, approximately 5.7 km^2^, corresponds to areas covered with vegetation (natural vegetation, gardens), bare soil, rocky outcrops, and beaches. Much of these areas are heavily degraded, often altered for centuries, with the exception of the cliffs and some beaches (Liarte, 2022).

### Fieldwork

Fieldwork was carried out between December 2023 and June 2025, with the aim, among others, of confirming the possible presence of the North African gerbil (*Gerbillus campestris*) in Melilla. For this purpose, 56 camera trap stations were installed, each equipped with a camera trap (Burton et al., 2015). The camera traps (Coolife, model PH700A) were active in video mode (15 seconds) with a 1-minute delay between each video, 24 hours a day for an average of 34 days. The sampling effort was therefore 1904 camera days. The traps were baited with a mixture of nuts and pieces of assorted fresh fruit. Given the recurrence observed during sampling throughout the camera trap operation period and the impossibility of independently identifying the detected individuals, only one record per location or camera trap station was considered. Each camera was fitted with a small triangular tunnel, 50 cm long by 15 cm wide (capture tunnel), designed and built by our own team. The bait was placed at the opposite end of the tunnel from the camera. The purpose of these tunnels is to bring the animal closer to the camera and keep it centred and in focus for as long as possible (Image 1).

**Image 1.**
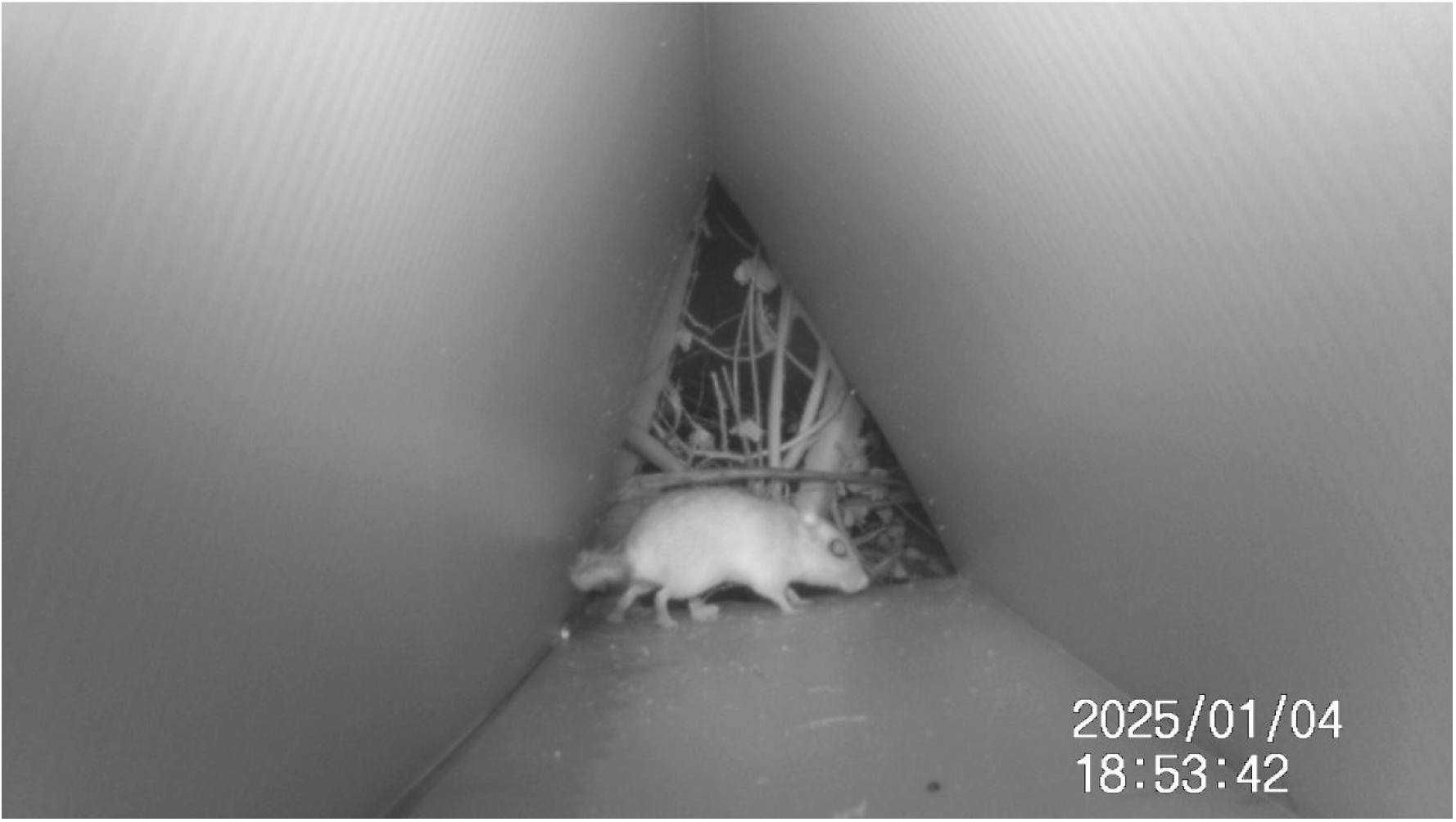
Specimen of Gerbillus campestris caught on camera inside the capture tunnel during fieldwork.

### Genetic Analysis of the Samples

For the taxonomic identification of the North African Gerbil, epithelial tissue samples were collected from the ear of one specimen, along with faecal samples. DNA was extracted using the InnuPREP DNA Mini Kit 2.0 (IST Innuscreen GmbH, Germany) according to the manufacturer’s protocol. Mitochondrial cytochrome b (*CYTB*) was amplified and sequenced using the following set of primers (5’-3’): L14723 [ACCAATGACATGAAAAATCATCGTT] and H15915 [TCTCCATTTCTGGTTTACAAGAC]. The *CYTB* was amplified using the following protocol: an initial denaturation at 95 °C for 15 min; 35 cycles of 95 °C for 30 s, 50 °C for 20 s, and 72 °C for 60 s; and a final extension at 72 °C for 5 min. The PCR product was sequenced using the Sanger method. Extracted DNA (40 μL) was used for Ultra-conserved elements (UCE) sequencing. Sequencing was performed in RAPiD Genomics (Gainesville, FL, USA).

### Variable estimation

Coverage: The total cover value (tree, shrub, and herbaceous) was estimated visually, using ranges of 0-25%, 25-50%, 50-75%, or 75-100%, always by the same researchers with over 15 years of experience in similar work.

### Anthropogenic impact: Three levels were established

High: presence of infrastructure (roads, reservoirs, housing, etc.) or human activities (crops, quarries, other industries, street markets, etc.) within a radius of 500 m or less.

Medium: presence of infrastructure (roads, reservoirs, housing, etc.) or human activities (crops, quarries, other industries, street markets, etc.) within a radius of 500 to 1000 m.

Low: presence within a radius greater than 1000 m of infrastructure (roads, reservoirs, housing, etc.) or human activities (crops, quarries, other types of industries, street markets, etc.).

## Results

Of the 56 camera trap stations installed, the presence of *Gerbillus campestris* was detected in 6 of them, representing a capture success rate of 10.71%. Some variables collected in the habitats where they were located are shown in Table 1.

**Table 1.** UTM (X; Y) locations of the camera traps where Gerbilluis campestris appeared; Alt: Altitude in meters; EVD: Dominant plant species; Cob: Total vegetation cover and Imp. Antro: Anthropogenic impact. The specimen marked in green is the one from which tissue and faecal samples were taken and on which body measurements were performed.

| X | Y | Alt (m) | Habitat | EVD | Cob (%) | Imp. Antro |
| --- | --- | --- | --- | --- | --- | --- |
| 504779 | 3907703 |  | Pastizal | *Salsola oppositifolia* y *Pinus halepensis* | 25-50 % | Alto |
| 504710 | 3907279 | 87 | Pastizal | *Salsola oppositifolia* | 25-50% | Alto |
| 504678 | 3907355 | 91 | Pastizal | *Salsola oppositifolia* | 0-25 % | Alto |
| 504734 | 3907271 | 86 | Pastizal | *Ligum spartium*, *Thymelaea hirsuta* | 50-75% | Alto |
| 503136 | 3906659 | 73 | Matorral | *Pistacea lentiscus*, *P. halepensis*, *Whitania frutescens* | 75-100% | Medio |
| 503073 | 3906330 | 24 | Bosque | *Robinia pseudoacacia* | 75-100% | Medio |

At one of the stations where the species appeared, a cage trap was set to capture a live specimen. The capture took place 24 hours after the cage trap was set. The body measurements taken of the captured specimen are shown in Table 2.

**Table 2.** Body measurements of the captured specimen. Cc: head-body measurements in cm; C: tail measurement in cm; Lpt: hindquarter length in cm and P: body weight in grams.

| Species | Locality | Cc | C | Lpt | P |
| --- | --- | --- | --- | --- | --- |
| *Gerbillus campestris* | Barranco del Nano | 8,5 | 13,5 | 2,5 | 37 |

### Genetic analysis results

Comparison with sequences available in GenBank revealed that the obtained *CYTB* sequence clusters with those of *Gerbillus campestris*. Analysis of the UCE loci placed the newly obtained sample within the *Gerbillus campestris* clade, clustering with previously identified individuals from our database. These results confirm the taxonomic identity of the specimen and validate the occurrence of *Gerbillus campestris* at the newly reported locality.

## Discussion

Ángel Cabrera, in his exhaustive works on Spanish (Cabrera, 1914) and Moroccan (Cabrera, 1932) mammals, makes no mention of any species of the genus *Gerbillus* in Melilla, nor, of course, in the Iberian Peninsula. This absence is significant given the precision and breadth of the work carried out by the Spanish zoologist at the beginning of the 20th century. Even before Cabrera, the species had not been recorded, to our knowledge, in what is now the Spanish

### Autonomous City of Melilla

The earliest possible record of this species appears as recently as 1986 (Aulagnier & Thevenot, 1986). In this study, as in the 2017 study (Aulagnier et al., 2017), the authors cite *Gerbillus campestris* as present in a grid square encompassing Melilla (approximately 12 km^2^) and a large area of Moroccan territory. In the first case, 0.5 x 0.5 degree grid squares (approximately 50 km per side) were used (Aulagnier & Thevenot 1986), and in the second, 0.25 x 0.25 degree grid squares (approximately 25 km per side) (Aulagnier et al., 2017). Both studies cover a considerable area of land beyond Spain’s borders. Neither study explicitly confirms the species’ presence in Melilla. Both projects were also carried out exclusively in Moroccan territory.

Closely related to the work of Aulagnier & Thevenot in 1986 are the citations by Palomo & Gisbert, 2002 and Palomo et al., 2007, whose references to the presence of *Gerbillus* (*Dipodillus*) *campestris* in Melilla rely on the former. Therefore, it is not possible to validate them because both citations are based on a previous, unverified bibliographic reference.

For their part, Yus et al., (2013) cite the species. However, this is a publication of a popular nature, which, as in the previous cases, references a citation copied from earlier, unconfirmed citations.

The latest review of mammals in Melilla also rules out the possibility of any valid records for the species (Román et al., 2025). Despite the relative abundance of different records of the species in Melilla, 5 in total (but three if the same Aulagnier & Thevenot, 1986 and Aulagnier et al., 2017 are considered, and likewise those of Palomo & Gisbert, 2002 and Palomo et al., 2007), in no case has it been possible to verify the existence of a verifiable record that confirms them (Román et al., 2025).

We have only found one record of the genus *Gerbillus*, specifically *G. gerbillus*, that could be considered valid for Melilla. This record, made in the Río de Oro in 1902, describes 5 males and one female (Riggenbach, 1903) and was collected and translated by Ángel Cabrera in 1904 (Cabrera, 1904). Based on Cabrera’s own comments, we must assume that the records were made in the Spanish section of the river: “Mr. Font submitted several descriptions of new vertebrate species from the Río de Oro, collected by F. W. Riggenbach and described by various specialists in foreign publications (…) Since it is advisable that descriptions of new species from Spanish territories be published in our Society even if they have already appeared in other journals, it was agreed that they be reproduced in the BULLETIN.” Therefore, this record is the first scientifically valid one that confirms the presence of *Gerbillus campestris* in Melilla and, consequently, in Spain and Europe.

## Acknowledgments

This study was funded by the Directorate General for Sustainability of the Department of Environment and Nature of the Autonomous City of Melilla, under contract number 2024056588. We appreciate the collaboration and helpfulness of Paola Calzado Liarte throughout the project.

